# ZNF638 represses the transcription of HBV closed circular DNA involving HUSH complex-mediated histone modifications of epigenetic silencing

**DOI:** 10.1101/2025.05.15.654218

**Authors:** Sifan Meng, Xiaowei Sha, Guoyan Zhang, Binwei Duan, Jing Wang, Yang Si, Feng Li, Qiong Wu, Shan Cheng, Wei Ding

## Abstract

Hepatitis B virus (HBV) infection poses a critical threat to public health burden worldwide. As the major template of viral transcription, the covalently closed circular DNAs (cccDNAs) of HBV was known to form microchromosomes and interact with various host proteins. In the present study, ZNF638 was found to significantly reduce the transcription of HBV into pgRNAs, and subsequently reduce cccDNA formation and viral propagation. We discovered that ZNF638 was able to bind the *preS* and *S* gene regions of HBV, repressing the transcription of cccDNA upon increased modification of H3K9me3 mediated by SETDB1. This epigenetic silencing of cccDNA transcription was further demonstrated to involve the activity of HUSH, a multi-protein complex to mediate epigenetic modifications of histone into suppressed nucleosomes. In *in vivo* models, we evaluated the effects of ZNF638 deficiency on HBV-targeting siRNA therapy. A significant reduction of the efficacy was observed in animals with reduced ZNF638 expression. In summary, we report that ZNF638 is a potent host factor to repress the transcription of HBV DNAs and to attenuate HBV infection. The role of ZNF638 on HBV repression involves its binding to cccDNA at specific regions for recruiting HUSH complex to write H3K9me3 histone marks for epigenetic silencing. ZNF638 serves as a critical host factor to associate HBV infection and disease progression. Additionally, ZNF638 serves as a potential indicator to predict the outcome of HBV-targeting siRNA therapy.

## Introduction

Hepatitis B virus (HBV) infection and its associated diseases remained to be a significant public health burden worldwide, especially in eastern Asian countries. The outcome of HBV infection is determined by complex host and viral factors, and the clinical manifestation of HBV infection can be observed as acute, chronic, or occult cases. Chronic hepatitis B (CHB) affects over 290 million people globally, whom mostly are of higher risk of developing fibrosis, cirrhosis, liver failure and hepatocellular carcinoma (HCC)^1^. The treatment of CHB is focused to reduce the risk of liver disease progression and prevent its harmful clinical sequelae, whom ideally to eradicate viral infections or at least partially by constantly inhibit viral replication^2, 3^. Many of the current anti-HBV therapeutics did show beneficiary from control of disease progression, nonetheless, lifelong treatment and surveillance are recommended for the majority of patients, as viral reactivation often occurs after medication cessation and drug resistance, which frequently developed during the treatment process^4, 5^. The major causes leading to the failure in CHB treatments is the ineradicable existence of covalently closed circular DNA (cccDNA) of HBV, which is a stable episomal form of the virus genome decorated with host histones and non-histone proteins^6^.

HBV is a small enveloped DNA virus carries a partially double stranded relaxed circular (rc) genome of 3.2 kb^7^. HBV virions attach to heparansulfate proteoglycans (HSPG), and then bind to the specific at high-affinity viral receptor (i.e., sodium taurocholate cotransporting polypeptide (NTCP) on the surface of hepatocytes^8^. After uncoating, the rcDNA in the HBV nucleocapsid is transported into the nuclei and converted into forms of cccDNA microchromosomes. Using cccDNA as the transcription template, HBV RNAs in various forms were produced by the host RNA polymerase II, including 3.5 kb pregenomic RNA (pgRNA) encoding viral polymerase (pol) and HBV core protein (HBc), 3.5 kb preCore mRNA for translating e-antigen (HBeAg), 2.4/2.1 kb *preS/S* mRNA encoding large (L), medium (M), and small (S) HBsAg respectively, as well as a 0.7 kb mRNA encoding regulatory protein HBx^9^. In the cytoplasm, HBc proteins encapsulate the pol and pgRNAs into nucleocapsid, where pgRNA is reverse transcribed into rcDNA by pol. The capsid carrying rcDNAs can be coated by viral envelope for virion egress, or redirected to the nucleus to replenish the cccDNA pool^10–12^. Accumulating evidence suggested that epigenetic modifications of cccDNA are functionally important to viral replication and define the outcome of chronic HBV infection^13^. However, the underlying mechanism for cccDNA microchromosome formation and modulation of its transcriptional activity in hepatocytes is not well understood.

As an important eukaryotic cellular defense mechanism to resist the introduction of foreign “non self” genetic material from infectious agents, the human silencing hub (HUSH) complex has been characterized as a multi protein assembly of three proteins, MPP8, TASOR, and PPHLN1^14^. Studies have shown that HUSH complex is involved in the silencing of integrated HIV-1 and unintegrated viral DNA AAV (adeno-associated virus)^15, 16^. RNA-mediated HUSH recruitment appears to be a common process for determining many HUSH targets (LINE1, KRAB-ZFP genes)^17, 18^. The recognition of nascent transcripts by Periphilin and their binding to target gene loci is a key step in initiating HUSH repression. The anchoring of HUSH to target loci subsequently recruits SETDB1 and resulted in histone H3K9me3 deposition^19, 20^. Recently, the recruitment of HUSH was reported through certain DNA-binding nuclear protein, such as ZNF638, also known as NP220, which exhibits sequence specificity in cases of unintegrated viral infections of murine leukemia virus (MLV) and recombinant adeno-associated virus (AAV)^21, 22^. Such noncanonical HUSH-mediated silencing of the transgenes was suggested to require the DNA-binding activity of ZNF638 to interact a functional chromodomain of MPP8, which was different to what has been identified for L1 silencing of retroviral integrations. The activation of SETDB1 methyltransferase to deposit repressive histone marks of H3K9me3 was reported to as a subsequent event following the recruitment of HUSH components upon the interaction of ZNF638 with unintegrated MLV genome.

Prior to the conversion into cccDNAs, rcDNAs released from the HBV capsids are transported into nuclei and undergo a series of DNA repair processes, during which deproteinated rcDNA (DP-rcDNA) and closed minus-strand rcDNA (CM-rcDNA) can be identified as putative transitional intermediates in the process of rcDNA-to-cccDNA conversion. The accumulated cccDNA from replication in the nucleus associates with histones and other non-histone proteins to form microchromosome^23^. In HBV producing hepatic cell lines, the HBV genome is integrated into host chromosomes in a defined low copy number, much less as compared to the amount of replicating cccDNAs. These HBV cell models (HepAD38 and HepG2.2.15) were used to study the epigenetic modulation viral transcription where many histone markers on the HBV cccDNA microchromosome were identified. Therefore, we postulate that depletion and reconstitution of ZNF638 expression in these cell models prior to HBV infections will provide important information for evaluating the role of HUSH in epigenetic silencing of HBV transcription.

In the present study, we investigated whether ZNF638 played an important role in HBV infection, focusing on the regulation of HBV cccDNA transcription. We found ZNF638 was able to repress the transcription of HBV cccDNAs upon its specific binding at the *preS* and *S* regions. As the methylation of histones decorated the cccDNA microchromosomes was known to be responsible for it silencing in transcription, we determined that SETDB1, a key histone methyltransferase for writing H3K9me3 marks, was required for ZNF638 repression on HBV transcription. Since SETDB1 was reported to involve HUSH mediated retroviral transgene expression silencing, we explored whether HUSH complex also participated to suppress HBV infection in context of inhibiting cccDNA transcription with epigenetic modifications. We found that the components of HUSH complex indeed contributed in SETDB1 dependent H3K9me3 for the silencing of the cccDNA transcription, where the binding of ZNF638 to cccDNAs appeared to be a prerequisite condition. Lastly, we tested whether the cellular ZNF638 level was a critical factor to influence the efficacy of HBV-targeting siRNA therapy, we demonstrated that ZNF638 deficiency compromised the functional inhibition of HBV infection following RNAi (RNA interference) mediated inhibition of candidate drug treatments, indicating the importance of ZNF638 and its value for the evaluation of epigenetic silencing in connection with the application in siRNA based anti-HBV therapy.

## Results

### Upregulated ZNF638 expression is associated with active HBV replication

The previous findings of ZNF638 for its intriguing roles in viral infections, such as MLV and AAV^21, 22^, prompted us to characterize the expression dynamics of ZNF638 during the course of HBV infection using cccDNA-producing *in vitro* cell models (Fig 1A). HepAD38 is a cell line that has an integrated HBV genome and can produce infectious viral particles inducibly under tetracycline-withdrawal culture conditions. HepG2-NTCP cells, derived from HepG2 background and reconstituted HBV entry receptor NTCP, serves as a convenient *in vitro* model and widely used for HBV laboratory studies. The levels of HBV DNA (rcDNA and intermediates generated during the cccDNA repair process, for example, DP-rcDNA and CM-rcDNA) were determined in native HepG2-NTCP cells following HBV infections (Fig 1B). We found that ZNF638 expression was upregulated significantly as detected at both mRNA and protein levels post HBV infections (Fig 1C). The cell viability was not significantly affected at the HBV infection doses applied in the experiment (Supplementary Fig 1A, B). To investigate whether ZNF638 is functionally involved in HBV infection, we deployed CRISPR technology in HepG2-NTCP (Supplementary Fig 1C) to construct a sgZNF638 cell pool, which was of an advantage for averaging out the potential off-target effects, for studies of HBV infection and transcription evaluation. We also established a ZNF638 overexpression model through transient transfection of a CMV-driven cassette in a piggyBac vector. Following infection of cells with HBV DNA for 6 days, we observed that HBV DNA levels were significantly reduced (Fig 1D) in ZNF638 overexpression HepG2-NTCP cells (ZNF638oe); whereas increased in sgZNF638 cells (Fig 1E). Additional results from FISH assays verified with increased signals of HBV DNA staining in sgZNF638 cells under the same infection conditions (Fig 1F). The results suggested that ZNF638 functioned to inhibit HBV production. In HBV-producing HepAD38 cells, higher levels of ZNF638 were observed in tet-off induction conditions (Fig 1G, H), in positive correlation with active HBV production. We explored the effect of either sgZNF638 or ZNF638oe on HBV DNA levels in HepAD38 cells as well. We found that altering ZNF638 expression affected the HBV DNA levels in both sgZNF638 and ZNF638oe HepAD38 cells (Fig 1I, J), especially in HepAD38 ZNF638oe cells where viral production was significantly attenuated (Fig 1I). We conducted similar experiments in an alternative cell model of HepG2.2.15, derived from human hepatoma HepG2 cells and integrated a full HBV genome. Similar results and conclusions were obtained as demonstrated in Supplementary Fig 2.

**Fig 1.**
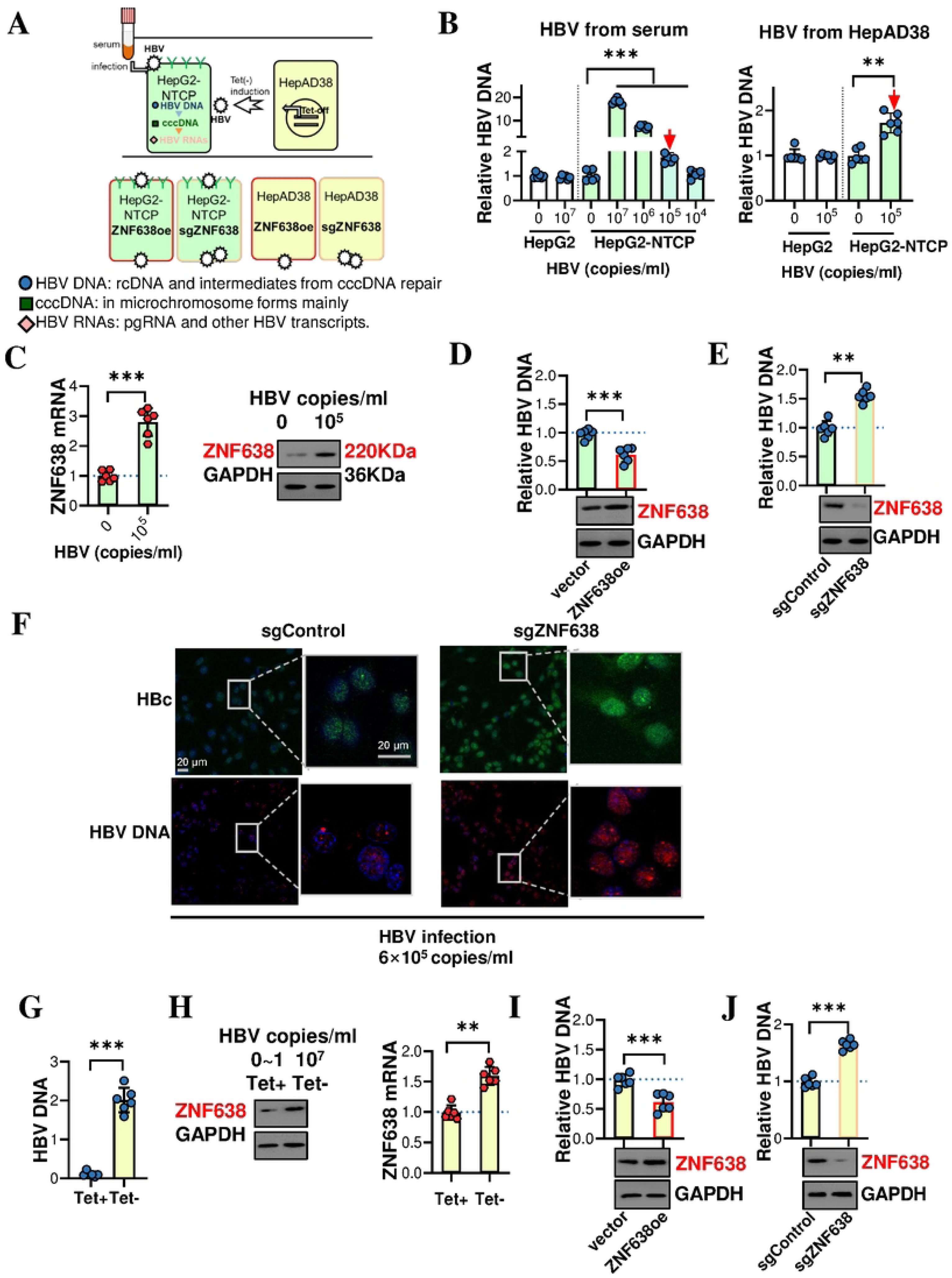
Correlational investigation of ZNF638 levels with HBV replication in hepatocyte cell lines. **(A).** *In vitro* laboratory models for HBV infection and production used for the present study. HepG2-NTCP (NTCP-overexpressing) cells were used for HBV infection experiments; HepAD38 (HBV integrated, tet-off inducible) cells were mainly used for HBV production and cccDNA preparation. Both cell lines were subjected to ZNF638 was overexpressed *via* piggyBac transfection or gene knocked out with CRISPR/Cas9 to construct *in vitro* models in experiments. (**B).** HBV DNA levels (rcDNA and intermediates during cccDNA repair process) in HepG2-NTCP cells (HBV infectable) and HepG2 cells (lacking NTCP for HBV infection) infected with either HBV prepared from patient serum at increasing doses (6× 10^4^, 10^5^, 10^6^ and 10^7^ copies/ml) for 96 h, or HBV produced from HepAD38 culture (6×10^5^ copies/ml) for 96 h. **(C).** ZNF638 expression in 6×10^5^ copies/ml HBV infected HepG2-NTCP cells at 6 d post infection as detected by RT-qPCR and western blot. **(D-E).** HBV DNA levels at 96 h post HBV infection (6×10⁵ copies/ml) in ZNF638-overexpressing HepG2-NTCP cells (ZNF638oe) at 48 h post-transfection, or in HepG2-NTCP stable pool (sgZNF638) prepared from CRSPR editing. (**F).** HBc IF or HBV DNA FISH in infected sgZNF638 HepG2-NTCP cells. SgZNF638 HepG2-NTCP cells were infected with HBV at 6×10⁵ copies/ml for 6 days. Images were collected under a 63× objective. (**G).** HBV DNA levels in HepAD38 cells at day 6 with or without the presence of 0.3 mg/ml tetracycline. **(H).** ZNF638 levels in HepAD38 cells determined by western blot and RT-qPCR under conditions as in **(G)**. **(I-J).** HBV DNAs in piggyBac-ZNF638 transfected HepAD38 cells (ZNF638oe), or in sgZNF638 edited HepAD38 pool, following Tet release of 6 days to induce HBV replication. ND: not detected.

### ZNF638 represses the transcription of HBV cccDNA

As the replication of HBV in host cells required cccDNA as the template of transcription, we hypothesized that the inhibitory effect of ZNF638 involved the suppression on cccDNA transcription. The production of HBV virion in sgZNF638 and ZNF638oe cell models were investigated with the quantitative detection of HBV cccDNA, its converted product rcDNA and transcribed pgRNA from rcDNA, together with the e antigen (HBeAg) as a translated protein product of HBV RNA. Knockout of ZNF638 in HepG2-NTCP cells markedly increased the HBV virion production, as evidenced by elevated levels of HBeAg with increased accumulation of HBV DNAs, transcribed pgRNAs (Fig 2A, B) and other HBV RNA species (Supplementary Fig 3A). In ZNF638oe HepG2-NTCP cells, the results were completely opposite (Fig 2C and Supplementary Fig 3B), where all the detected molecules were significantly decreased. In HepAD38 models, the effects of ZNF638 expression on HBV measures (Fig 2D, E and Supplementary Fig 3C, D) were consistent with the findings in HepG2-NTCP models. Similar results were obtained from HepG2.2.15 cells where the levels of ZNF638 were modulated by transient transfection (Supplementary Fig 4).

**Fig 2.**
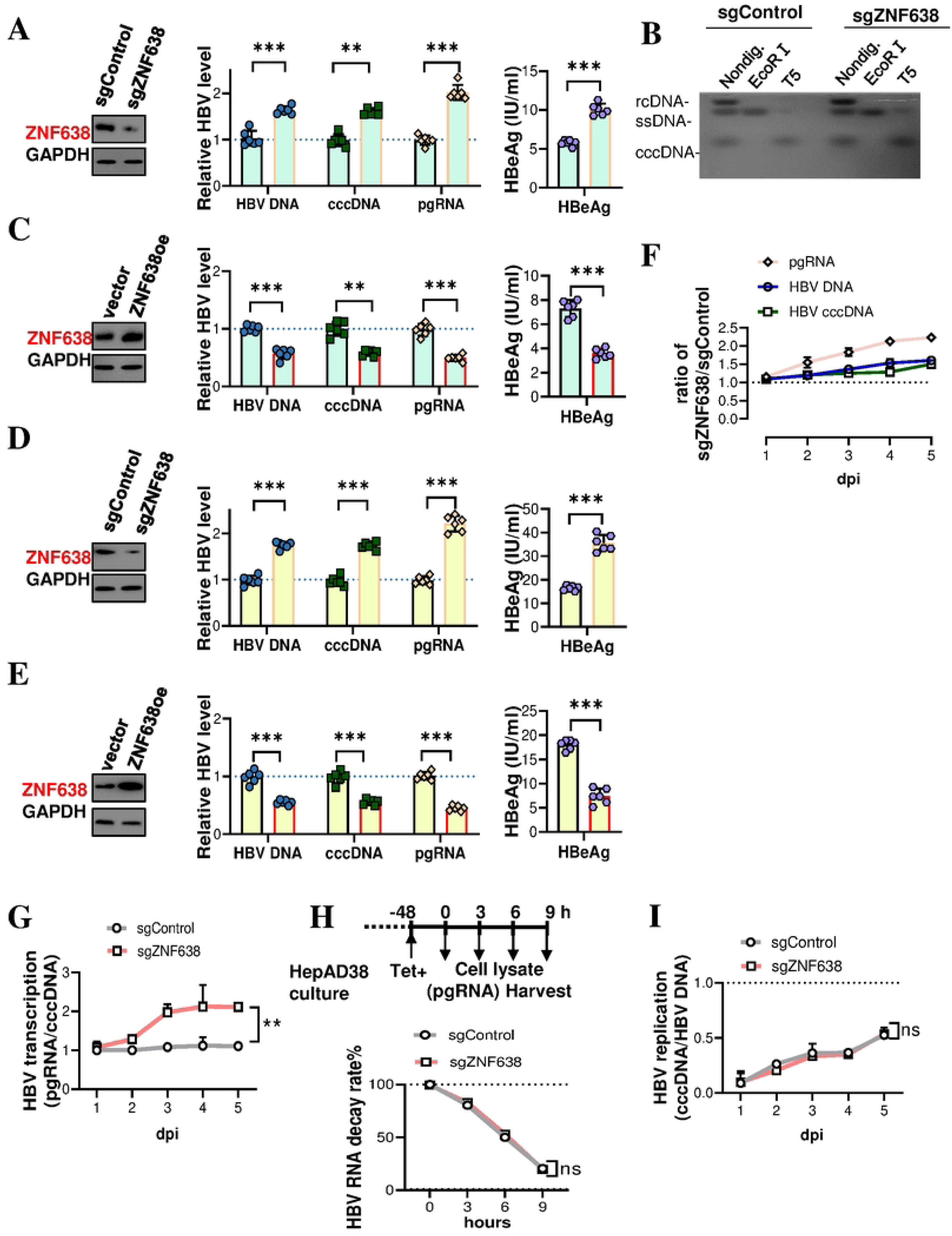
ZNF638 represses HBV cccDNA transcription. sgZNF638 **(A)** or ZNF638oe **(C)** HepG2-NTCP cells were infected with 6×10^5^ copies/ml HBV at 6 d post infections. HBV DNA (enriched in rcDNA), cccDNA (rcDNA, ssDNA and dsDNA cleared HBV DNAs from T5 exonuclease digestion), pgRNA, and e antigen were quantified using qPCR, RT-qPCR, ELISA or southern blot, respectively. **(B).** Southern blot analysis of HBV cccDNA in sgControl and sgZNF638-knockdown HepG2-NTCP cells at 6 days post-HBV infection (Nondig.: Non-digested DNAs). In sgZNF638 **(D)** or ZNF638oe **(E)** HepAD38 cells in Tet-free medium at 6 d for HBV induction, the levels of HBV DNA, cccDNA, pgRNA and e antigen were detected. **(F).** Kinetic quantification of HBV pgRNA and cccDNA production comparing in HBV-infected native HepG2-NTCP cells and ZNF638 knockout cells. **(G).** Transcription of cccDNA in HBV infected sgZNF638 HepG2-NTCP cells present as ratios of pgRNA to cccDNA. **(H).** Estimation of HBV RNA decay from normalized remaining percentage. Control or sgZNF638 HepAD38 cells were seeded at 48 h prior to the addition of tetracycline (time 0). The HBV RNA levels were determined from harvested cells at 0, 3, 6, 9 h. **(I).** Levels of cccDNA normalized to HBV DNA copies in HBV infected sgZNF638 HepG2-NTCP and compared to sgControl cells.

From results of all experimental conditions, the increased pgRNA levels were found in ZNF638 deficiency samples, which strongly suggested the transcriptional activity of cccDNA was repressed with the existence of ZNF638. The HBV transcription activity in ratios normalized to the controls, the production of pgRNAs was significantly reduced when ZNF638 were depleted during the time course of HBV infections (Fig 2F). When normalized the transcription products to template DNA abundance in cccDNA copy numbers, the inhibition of ZNF638 on cccDNA transcription appeared to be drastic as shown in Fig 2G. To exclude a possibility that ZNF638 might reduce the stability of pre-existed or newly transcribed HBV pgRNAs, we also performed experiments chasing for up to 9 hours in HepAD38 cells, in which no significant change was found (Fig 2H). However, the ratio of cccDNA over HBV DNA in both sgZNF638 and control samples exerted no significant changes (Fig 2I), which indicated that the changes in pgRNA were primarily from transcription regulation instead of the alterations in DNA template levels. Therefore, we concluded that the role of ZNF638 in HBV suppression was predominantly through the transcription inhibition of cccDNAs.

### ZNF638 suppresses the transcription of HBV rcccDNA without altering the repair of RrcDNA

To verify the inhibitory effect of ZNF638 on HBV cccDNA transcription, we employed the alternative recombinant systems for producing HBV cccDNA and rcDNA^24, 25^. In the prcccDNA plasmid, a chimeric intron and loxp sites were inserted into the HBV genome, and excised using Cre/loxP-mediated DNA recombination to form 3.3 kb rcccDNA. The chimeric intron can be spliced out from the RNA transcript, avoiding the disruption the translation of HBV proteins (Fig 3A). We cotransfected prcccDNA and pCMV-Cre plasmids into HepG2-NTCP cells of ZNF638 knockout or overexpression, and quantified levels of rcccDNA and pgRNA at 72 h post-transfection, when the pgRNAs were exclusively from the transcription of recombinant rcccDNAs. The activity of HBV transcription, as indicated in pgRNA to rcccDNA ratios, was significantly decreased in ZNF638 overexpression cells, whereas increased over the controls in sgZNF638 HepG2-NTCP cells (Fig 3B). Taking the advantage of the recombinant rcDNA system being used, the repairing of rcDNA could also be investigated. We annealed the HBV minus and plus sequences amplified separately by PCR, and then ligate a DNA flap and RNA-DNA oligo to produce RrcDNAs (Fig 3C). The levels of HBV DNA and cccDNA were quantified in RrcDNA transfected HepG2-NTCP cells of ZNF638 knockout or overexpression. No significant change was found in sgZNF638/ZNF638 overexpression cells and the controls as from the ratio of cccDNA to HBV DNA (Fig 3D). These results indicated that role of ZNF638 to inhibit the transcription of HBV cccDNA, but not to interfere the repair of rcDNA to produce cccDNA as the template.

**Fig 3.**
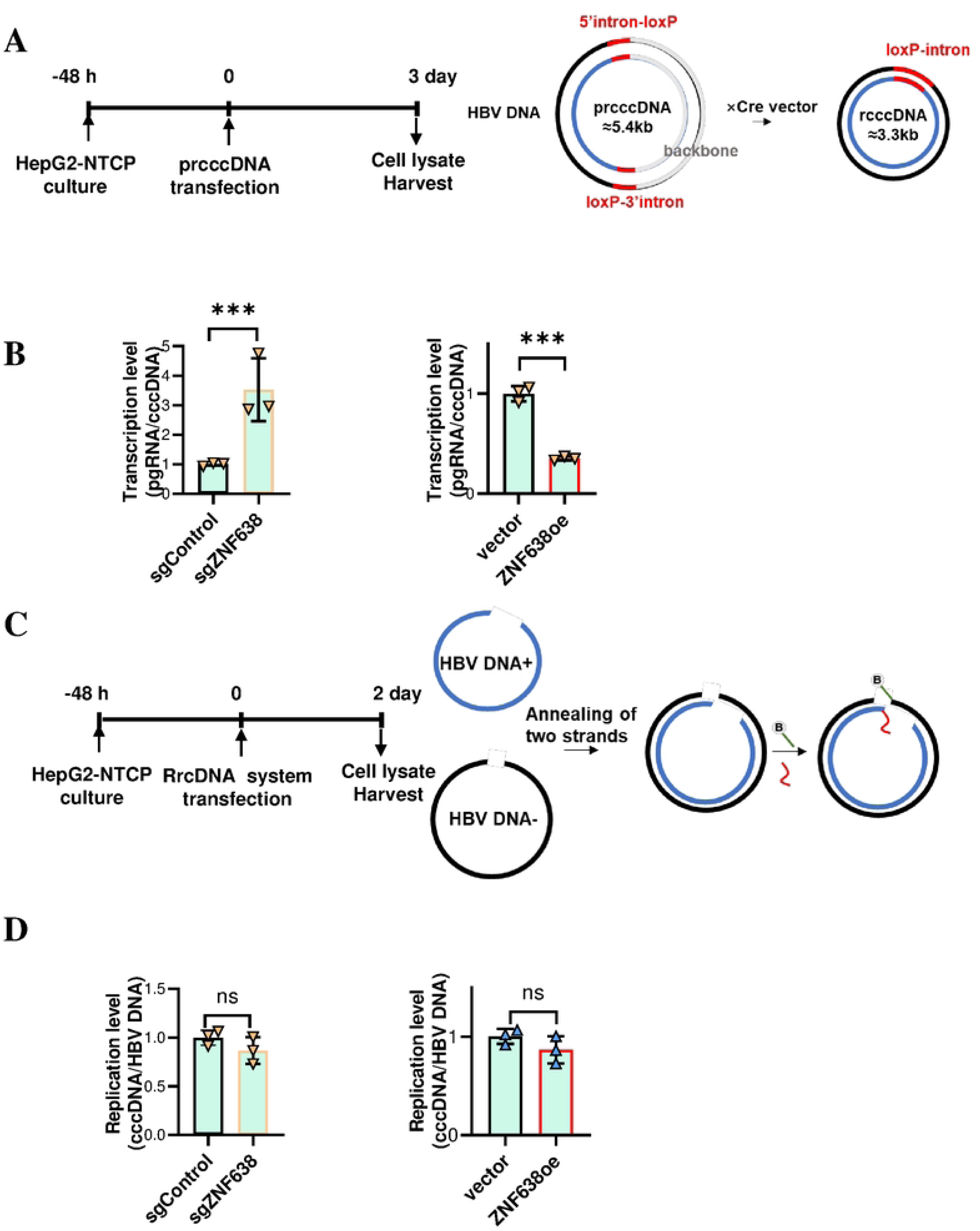
ZNF638 represses the transcription of HBV rcccDNA but does not affect the repair of HBV RrcDNA to cccDNA. **(A).** Principle of Cre/loxP-mediated recombination rcccDNA system. In the presence of Cre recombinase (provided in pCMV-Cre), the prcccDNA is excised at the loxP within a chimeric intron, where rcccDNA can be produced. The HBV RNA and cccDNA from ZNF638oe or sgZNF638 HepG2-NTCP cells were determined at 72 h following cotransfection of prcccDNA and pCMV-Cre (time 0). **(B).** The transcription of rcccDNA as in ratios of pgRNA to cccDNA in sgZNF638 or ZNF638oe HepG2-NTCP cells. **(C).** The production of recombinant HBV rcDNA (RrcDNA). HBV ssDNAs of either minus-strand (black) or plus-strand (blue) were prepared through PCR and annealed. The ligation with a DNA flap and RNA-DNA oligo was performed to form recombinant rcDNA (RrcDNA). The HBV DNA and cccDNA levels were determined at 48 h in ZNF638oe or sgZNF638 HepG2-NTCP cells following the transfection of HBV RrcDNA (time 0). Green line, biotinylated flap (b, biotin); Red line, RNA primer. **(D).** Normalized levels of cccDNA to HBV DNA copies in sgZNF638 or ZNF638oe HepG2-NTCP cells at 48 h post transfection of HBV RrcDNA.

### ZNF638 binds *PreS2* and *S* regions of HBV cccDNA

ChIP-PCR and ChIP-seq approaches were extensively used to identify the binding motifs of transcription factors, many of which belong to the zinc finger family. We isolated cccDNAs in native states from induced HepAD38 cells from volume cultures, then deployed incubation with an anti-ZNF638 polyclonal antibody for the magnetic pulldown enrichment. The retrieved sonicated DNA fragments were subjected to qPCR assays. A total of 20 pairs of primers covering the entire HBV genome were used for screening of positive signals. The normalized ChIP-qPCR results indicated the peak amplification covering the *preS2* and *S* gene regions (91-778 nt) (Fig 4A). We then conducted ChIP-seq experiments to characterize the ZNF638 binding of cccDNA. As shown in Fig 4B, the previous findings were verified and two peaks were identified at 91-318 nt (*preS2*) and 445-778 nt (*S gene*) regions, respectively. ChIP-qPCR assays were also carried out to calculate the fold enrichment of the identified peaks (Supplementary Fig 5). The binding of ZNF638 to HBV cccDNA was confirmed in extrachromosomal cccDNA isolated from and induced HepAD38 (Fig 4C) and HBV-infected HepG2-NTCP cells (Fig 4D). As we expected the binding of ZNF638 and HBV cccDNA could be manifested by the co-existence at cellular level, we performed FISH and IF staining in HepG2-NTCP cells infected with HBV. The results showed a clear colocalization of HBV cccDNA with ZNF638 protein within the nucleus (Fig 4E).

**Fig 4.**
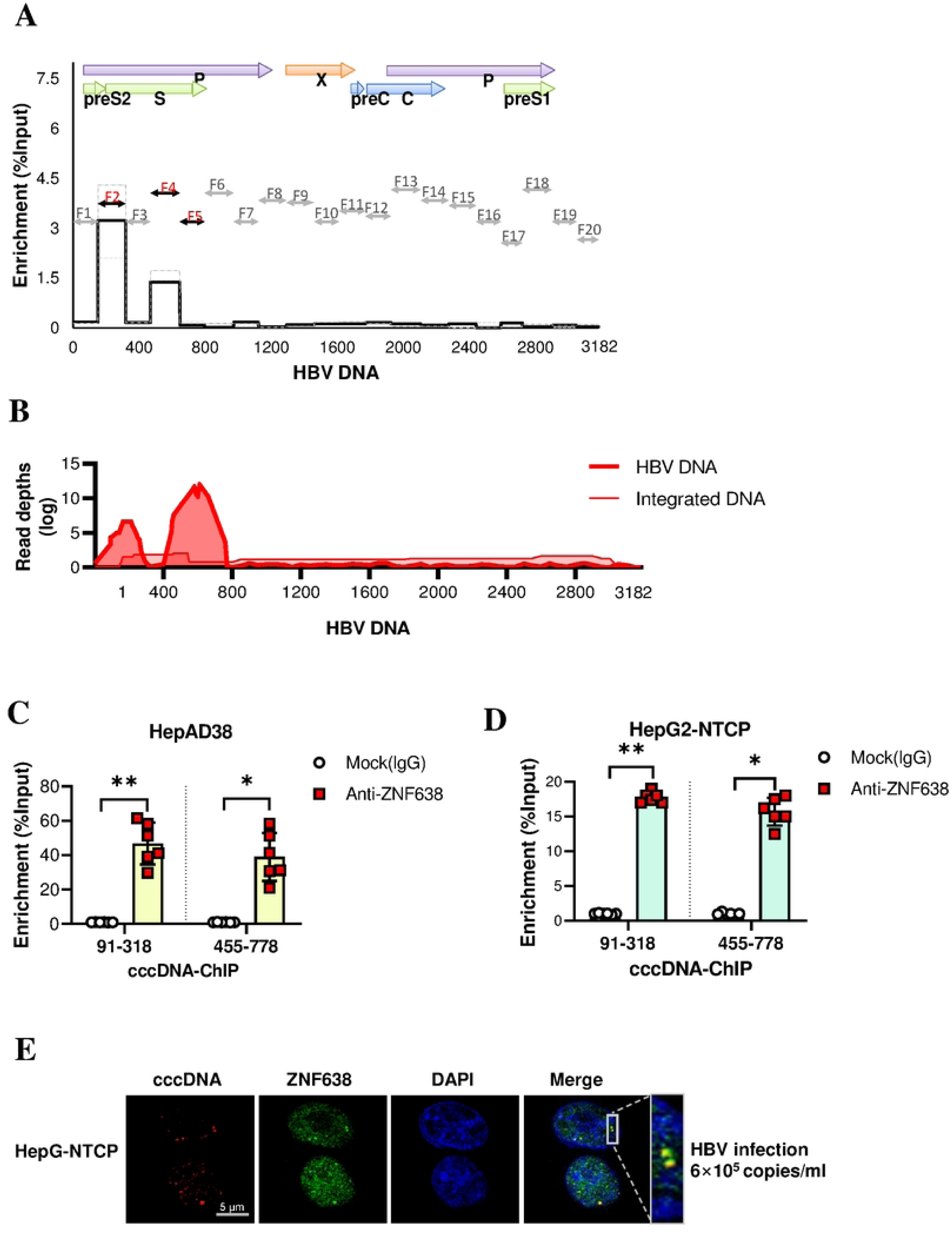
ZNF638 was able to bind directly to HBV cccDNA specific regions coding for *preS* and *S* genes. **(A).** The screening for ZNF638 binding regions across HBV genome by ChIP-qPCR using tiling primers in HBV producing HepAD38 cells. **(B).** ChIP-seq for ZNF638 binding peaks using DNAs isolated from HBV producing HepAD38 cells. **(C).** Binding of ZNF638 to cccDNAs (purified from HBV producing HepAD38 cells) as determined by ChIP-qPCR assays. **(D).** ChIP-qPCR assays to verify the binding of ZNF638 using cccDNAs purified from HBV infected HepG2-NTCP cells at 6 dpi. (**E).** Colocalization of HBV cccDNA and ZNF638 in HBV infected HepG2-NTCP cells. HepG2-NTCP cells were infected with HBV at a concentration of 6×10⁵ copies/ml for 6 days. The combined HBV cccDNA FISH and ZNF638 IF images were acquired at a 63× objective.

As HepAD38 cells carried genome integrated HBV sequences, which also contained the *preS2* and *S* gene regions, we questioned whether the identified ZNF638 binding peaks were artifacts from integrated viral DNAs. We analyzed the sequencing data for viral-human chimeric DNA reads, and found the counts were only about 20, merely equivalent to the input basal peak levels (25 and 62 as shown in Fig 4B and Supplementary Fig 6, in HepAD38 cells under both non-induced and induced conditions), whereas the ChIP-seq peak counts reached 2393 (Fig 4B), which was nearly 100-fold higher. Therefore, we believe that ZNF638 was indeed able to bind cccDNA directly at the identified specific regions. Meanwhile, we could not rule out the possibility that ZNF638 also bound to genome integrated HBV sequences, or the ZNF638 recognition site for genomic integrated HBV DNA copies might have different signatures.

### ZNF638 knockdown alters histone modifications on HBV cccDNA microchromosomes

Studies have shown that cccDNAs existed in forms nucleosomes decorated microchromosome, where histones were subject to various modifications. High abundances of H3K27ac, H3K4me3, and H3K9me3 modifications were observed, with H3K27ac^26^ and H3K4me3^27^ being associated with significant stimulation of transcription, whereas H3K9me3 was found to exhibit strong inhibitory activity on transcription^28^. Since the HBV induced ZNF638 was not able to completely stall viral replication as demonstrated in earlier results, we hypothesized that H3K9me3 was likely to involve in ZNF638 associated silencing on cccDNA transcription. We designed and conducted nuclear cccDNA pulldown assays (Fig 5A). Nuclear fractions were separated and then incubated with biotinylated oligonucleotides specific to HBV cccDNA (cccDNA bait). Streptavidin-coated magnetic beads were then used to precipitate the cccDNA-bait complex along with its associated proteins. We found that ZNF638 knockdown decreased cccDNA associated H3K9me3 in HepAD38 cells or HBV-infected HepG2-NTCP cells, but not H3K4me3 and H3K27ac (Fig 5B). The results demonstrated that the H3K9me3 suppression on cccDNA transcription required the participation of ZNF638. The data also suggested that the H3K27ac and H3K4me3 were not able to influence cccDNA transcription required ZNF638.

**Fig 5.**
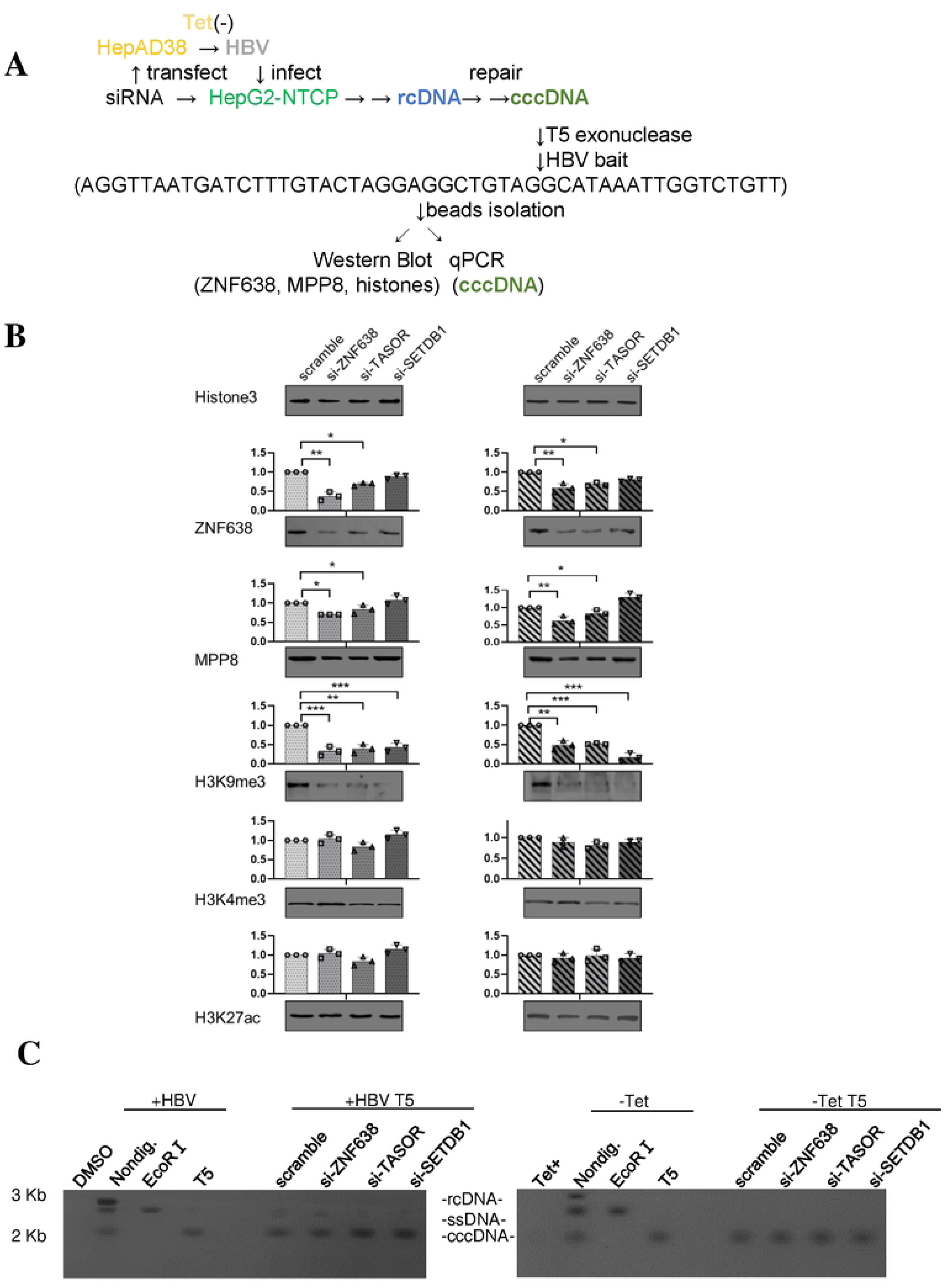
ZNF638 knockdown modulated histone modifications of HBV cccDNA microchromosomes. **(A).** Nuclear HBV cccDNA pulldown-WB procedures. After transfection with siRNAs targeting ZNF638, TASOR, and SETDB1 for 48 h, HepAD38 cells were transferred to Tet-free medium to induce HBV replication and cultured for an additional 6 days; HepG2-NTCP cells were infected with 6×10^5^ copies/ml HBV for 6 days. T5 exonuclease was used to digest isolated extracts to remove rcDNA, ssDNA and dsDNA. Following incubation with a specific biotinylated cccDNA bait, streptavidin-coated magnetic beads were employed to precipitate the baited cccDNA and associated proteins in the complex. **(B).** Western blotting for the detection of proteins of interest alone with various histone H3 in HepG2-NTCP cells (left) and HepAD38 cells (right). Semiquantitative measures from densitometry were normalized to controls and plotted as changes in folds (n=3). **(C).** Southern blotting for HBV cccDNA from HepG2-NTCP (left) and HepAD38 (right) cells. (Nondig.: Non-digested DNAs)

### Suppression of ZNF638 on HBV transcription requires SETDB1-mediated H3K9me3 of cccDNA microchromosome

Recent studies indicated that methyltransferase SETDB1 was a key enzyme to deposit repressive H3K9me3 histone marks during murine leukemia virus (MLV) infection and integration^21, 22^. We started to examine whether SETDB1 was necessary for ZNF638 on transcription silencing of cccDNA. In SETDB1 knockdown HepG2-NTCP cells, the levels of HBV DNA, cccDNA and pgRNA were significantly increased post infection (Fig 6A), and dose-dependently to the degrees of SETDB1 depletion (Fig 6B). From the ratio of pgRNA/cccDNA representing the activity of HBV transcription per templated, the significant increase in SETDB1 knockdown groups indicated that SETDB1-mediated H3K9me3 could be crucial for the regulation of cccDNA transcription at epigenetic levels (Fig 6C).

**Fig 6.**
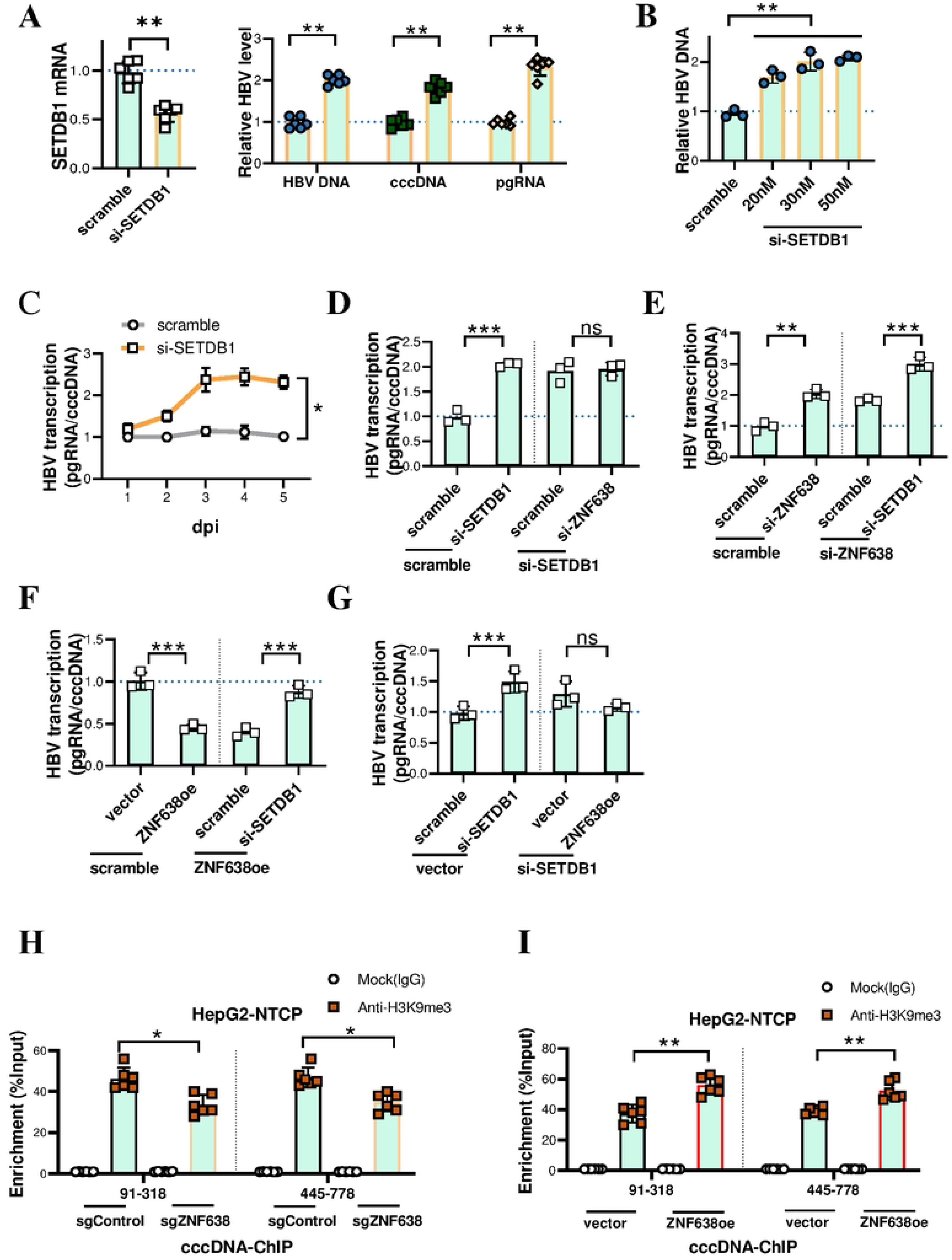
ZNF638 dependent epigenetic silencing of HBV cccDNA transcription required marking of SETDB1-mediated H3K9me3. **(A).** Levels of HBV DNA, cccDNA and pgRNA in SETDB1 knockdown HepG2-NTCP cells at 6 d of 6×10^5^ copies/ml HBV infections after SETDB1 siRNA transfection for 48 h. **(B).** Dose response of SETDB1 knockdown on increased HBV DNA levels. **(C).** Time course of HBV cccDNA transcription shown as the ratio of pgRNA to cccDNA for efficacy comparison. **(D-G).** Effect of SETDB1 siRNA transfection at 48 h on normalized HBV transcription levels at 6 d of 6×10⁵ copies/ml HBV infections in ZNF638 knockdown or overexpressing HepG2-NTCP cells. **(H-I).** ChIP-qPCR assays on cccDNAs for H3K9me3 enrichment at the ZNF638 binding regions using cccDNAs isolated from sgZNF638 or ZNF638oe HepG2-NTCP cells at 6 d post HBV infection.

Both of ZNF638 and SETDB1 were able to suppress cccDNA transcription into pgRNAs, the silencing capacity of SETDB1 appeared to be much more potent and dominant. In ZNF638 knockdown HepG2-NTCP cells, the enhancement of HBV transcription was detected upon additional depletion of SETDB1 (Fig 6D). In SETDB1 siRNA knockdown cells, reducing ZNF638 expression was not further increasing pgRNA levels, as the viral cccDNA transcription was nearly maximized (Fig 6E). In ZNF638oe models, the potency of SETDB1 was demonstrated with more significant results (Fig 6F, G). These results implied that SETDB1 functioned at the downstream of ZNF638 for the silencing of cccDNA. If ZNF638 is responsible for recruiting SETDB1, it is rational to predict that the H3K9me3 occupation sites will overlap with ZNF638 binding peaks, and alterations will be detected upon the changes in ZNF638 levels. ChIP-qPCR assays revealed that the enrichment of H3K9me3 at the ZNF638 binding regions of HBV cccDNA was decreased by sgZNF638 (Fig 6H) and enhanced by ZNF638 overexpression (Fig 6I). However, in SETDB1 knockdown cells, less inhibition of ZNF638 on HBV transcription was observed, although the binding of ZNF638 to cccDNA was not altered at the specific sites (Fig 5B). These results suggested that the binding of ZNF638 to cccDNA could facilitate the recruitment of SETDB1, which in turn deposited the repressive H3K9me3 histone modification, thereby inhibiting cccDNA transcription.

### ZNF638 mediated suppression of HBV cccDNA transcription is involved with HUSH complex

Although ZNF638 was able to facilitate SETDB1-mediated methylation of cccDNA histones, we proposed that it was not likely sufficient and that direct protein-protein interactions from their known structural domains might be involved. Alternatively, ZNF638 was previously discovered to bind HUSH complex; and reports demonstrated that HUSH complex mediated epigenetic silencing of viral integrated genes require SETDB1 activity on histone methylation. We hypothesized that the facilitation of SETDB1 recruitment to cccDNA was mediated by HUSH, which will help to explain why the H3K9me3 distribution matched ZNF638 peaks in cccDNA. We used RNAi technology to deplete each component of HUSH to evaluate their functional involvement in HBV cccDNA transcription in infected HepG2-NTCP cells (Fig 7A). The results showed that knockdown either of the three components of HUSH (MPP8, TASOR or PPHLN1) led to significant increase in HBV transcription with elevated levels of pgRNAs. ChIP-seq experiment was performed using a MPP8 antibody to investigate whether HUSH complex associated with cccDNAs. In HepAD38 cells, the most enrichment of MPP8 was found at *S* gene (445-778 nt), overlapping one of the ZNF638 binding sites (Fig 7B). The quantification of HBV cccDNA enrichment was derived from ChIP-qPCR (Fig 7C), where the weak interaction of HUSH with cccDNA at the *preS2* gene (91-318) region was verified. Nuclear cccDNA pulldown assays showed that the knockdown of ZNF638 decreased the recruitment of MPP8 on cccDNA (Fig 5). The ZNF638 occupation on cccDNA was also reduced by the knockdown of TASOR (Fig 5). The binding of MPP8 to the *S* gene position (445-778 nt) of HBV cccDNA was verified with extracellular cccDNA isolated from HBV infected HepG2-NTCP cells (Fig 7D). The knockdown of either ZNF638 or TASOR was shown to decrease SETDB1-mediated H3K9me3 (Fig 5). In ZNF638 knockdown HepG2-NTCP cells with MPP8 depletion, greater enhancement of HBV transcription was detected as compared to ZNF638 knockdown alone (Fig 7E). Similarly, in MPP8 knockdown cells with ZNF638 depletion, the cccDNA transcription was further enhanced following HBV infection (Fig 7F). In ZNF638oe HepG2-NTCP cells, knockdown MPP8 decreased the inhibitory effect on HBV transcription (Fig 7G). HBV transcription was observed increased in MPP8 knockdown cells. By overexpression ZNF638, the transcribed HBV RNA level was suppressed to normalized basal level and lower (Fig 7H). In sgZNF638 HepG2-NTCP cells, the probing for MPP8 binding at the HBV *S* gene positions decreased below control level in ChIP-qPCR (Fig 7I), whereas in ZNF638oe HepG2-NTCP cells, the enrichment was significantly increased (Fig 7J). Taken together, these results were in support of previous findings for the role of ZNF638 in retroviral infections, where the binding of ZNF638 increased the binding of HUSH complex to target loci and subsequent recruitment of SETDB1, causing the potent repression on the transcription of viral sequences. The deficiency in ZNF638 tampered the recruitment of HUSH at selected loci and significantly compromised the inhibition of cccDNA transcription.

**Fig 7.**
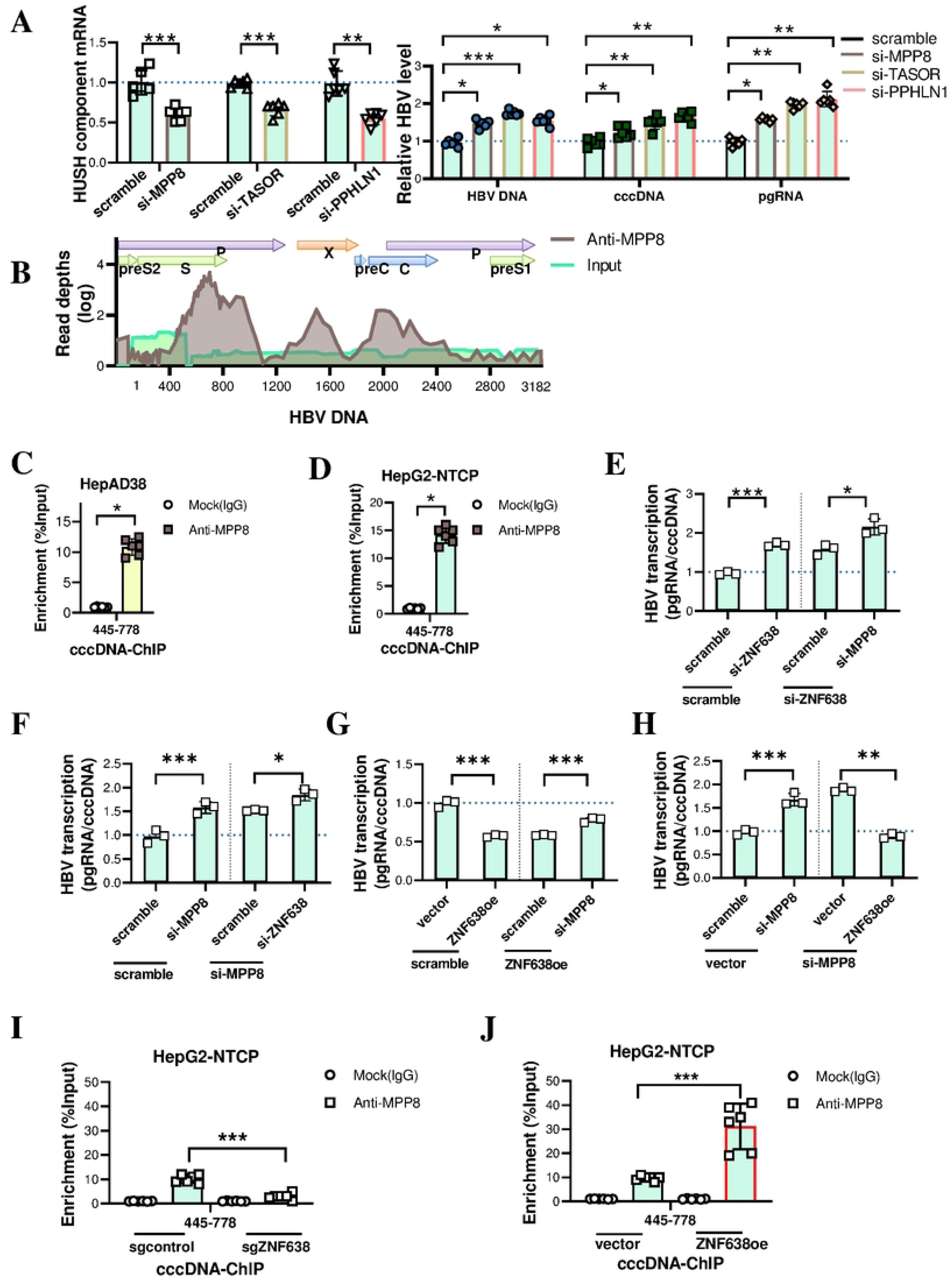
Knockdown individual component of the HUSH complex significantly attenuated the ZNF638 dependent inhibition on HBV transcription. **(A).** Knockdown levels of MPP8, TASOR or PPHLN1 from normalized RT-qPCR measures of mRNAs following siRNA transfection for 48 h in HepG2-NTCP cells (left); HBV DNA, cccDNA or pgRNA levels by qPCR or RT-qPCR at 6 d post HBV infections (right). **(B).** ChIP-seq of DNAs from HBV producing HepAD38 cells using an anti-MPP8 antibody for determining MPP8 binding sites over the HBV genome. **(C).** Validation of MPP8 binding at the *S* gene regions from ChIP-qPCR of cccDNAs isolated from HBV producing HepAD38 cells to obtain enrichment measures of statistical significance. **(D).** ChIP-qPCR of cccDNAs isolated from HBV infected HepG2-NTCP cells at 6 dpi for the quantification of MPP8 binding at the HBV *S* gene region. **(E-F).** MPP8 or/and ZNF638 siRNA transfected HepG2-NTCP cells and infected for 6 days, HBV transcription as determined by pgRNA production per cccDNA copies from RT-qPCR assays. **(G-H).** MPP8 siRNA or/and ZNF638 overexpression plasmid were used to transfect HepG2-NTCP cells, HBV transcription was assayed at 6 d post infection. **(I-J).** ChIP-qPCR assays in sgZNF638 or ZNF638oe HepG2-NTCP cells for enrichment of MPP8 binding to HBV *S* gene regions using cccDNA isolated at 6 d post infection.

### Deficiency of ZNF638 significantly compromised the efficacy of HBV-targeting siRNA therapy

Recently, anti-HBV siRNA drugs have developed for treating chronic hepatitis patients. GalNAc conjugated siRNA may also be applied to construct *in vivo* conditions of ZNF638 knockdown in the liver, allowing the evaluation of ZNF638 deficiency for its pathological significance during HBV infection. As earlier study suggested that AAV transduction could be influenced by HUSH complexes with the involvement of ZNF638. Due to this, the AAV-based HBV *in vivo* models were not considered in this study.

With the HBV transgenic animals of four experimental groups (Fig 8A), serum HBV DNA levels and secreted HBe antigens were assayed during the time course. ZNF638 siRNA administration significantly increased the serum levels of HBV DNA and HBeAg, and markedly reduced the efficacy of HBV siRNA treatment (Fig 8B). In ZNF638 knockdown mice, the liver HBV pgRNA and HBeAg increased to 286% and 151% respectively by the end of the experiment at day 29 (Fig 8C). HBV siRNA delivery was able to decrease liver levels of HBV DNA, pgRNA and HBeAg to 23%, 85% and 60%, however, ZNF638 siRNA administration alone it the values went up by 362% and 338% in pgRNA and HBeAg (Fig 8C). The mRNA expression of ZNF638 was also analyzed in the liver tissues of the mice. Results confirmed the siRNA efficiency of ZNF638 in the mice of ZNF638 siRNA group and co-administration group (Fig 8D). Neither organ toxicity nor weight loss in mice were recorded over the 29-day period treatment (Supplementary Fig 7). These supported our previous findings that active replication of HBV induced a responsive upregulation in ZNF638 expression, which subsequently acts to suppress HBV transcription in later infection phases, and might be significantly inhibit virus replication. However, deficiency of ZNF638 in host cells will cause infected HBVs rapidly enter and maintain in active life cycles. The *in vivo* data once again demonstrated that ZNF638 significantly inhibited the transcription of HBV RNA, the replication of viral DNA, and the production of antigen (HBeAg). As there is an argument based on findings that the transgenic mice model being used are not capable producing cccDNAs efficiently, implying the exact mechanism of ZNF638 function under the circumstance can be more sophisticated than that was discovered in cell line studies. Nonetheless, as a highly conserved nuclear factor, the importance of ZNF638 on HBV infection shall not be neglected. At least, the deficiency of ZNF638 significantly impaired the efficacy of HBV-targeting siRNA therapy, which deserves attention for clinical considerations.

**Fig 8.**
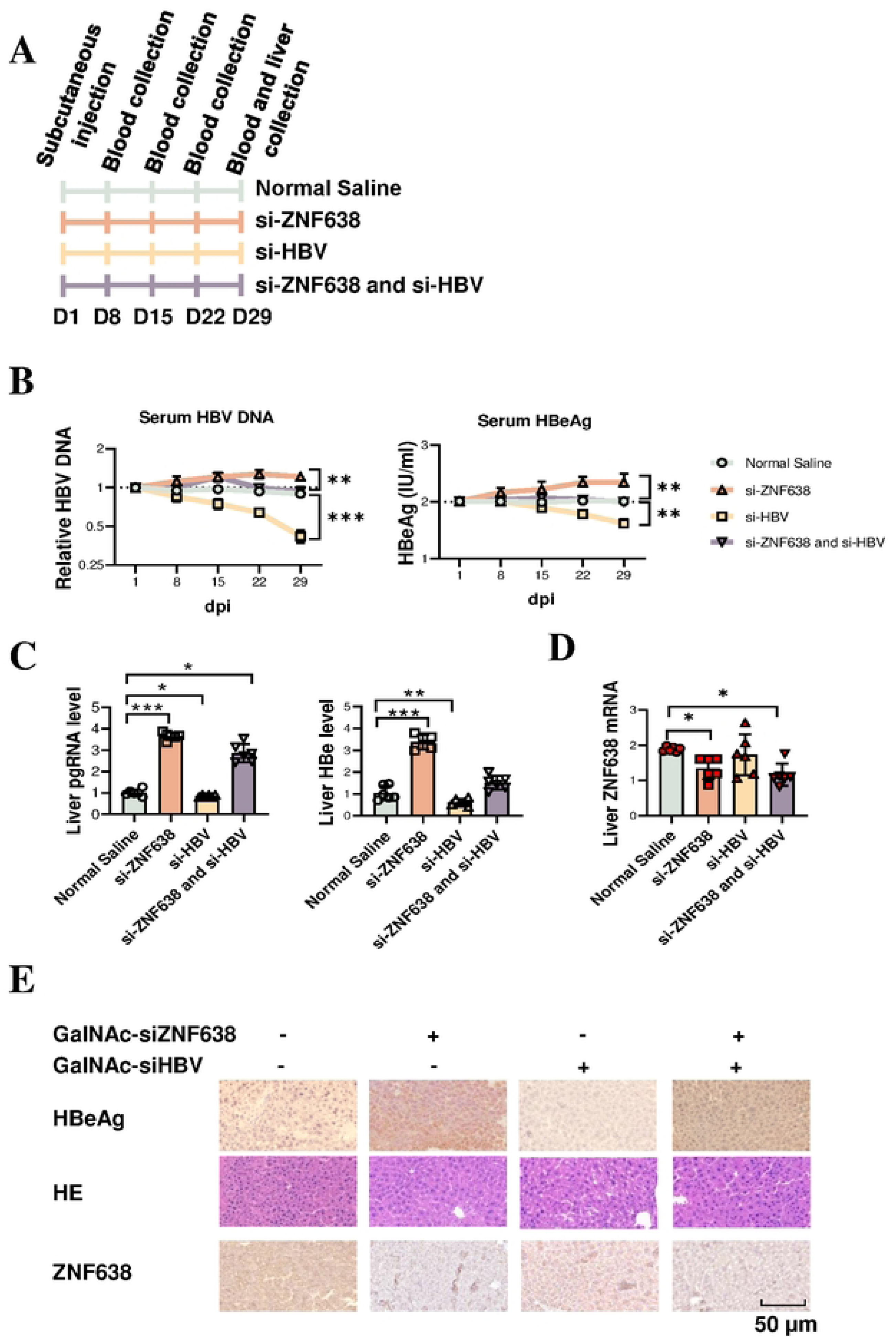
ZNF638 deficiency compromised the efficacy of HBV siRNA therapy. **(A).** HBV transgenic mice were randomized into each experimental groups according to serum levels of HBsAg and HBV DNA. A single dose of GalNac-siHBV (3 μg/g) and/or GalNac-siZNF638 (3 μg/g) were administrated through subcutaneous injections (day 0). Blood samples were collected from alternated lateral and medial canthus of the left or right eye alternately at day 8, 15, 22 and 29 (day sacrifice). **(B).** Serum levels of HBeAg and HBV-DNA as determined by ELISA and RT-qPCR. **(C).** Normalized tissue levels of pgRNA in the liver quantified by qPCR. **(D).** RT-qPCR for ZNF638 mRNA levels in hepatic tissues. **(E).** Immunohistochemistry and HE staining of collected liver tissue sections (40 x).

## Discussion

Based on the data from present study, we proposed a possible model illustrating the mechanism on the role of ZNF638 to suppress HBV transcription. The binding of ZNF638 to the *preS2* and *S* gene regions of HBV cccDNA, especially at the S gene region, is required for the recruitment of HUSH complex at the adjacent loci, then enables SETDB1 to write H3K9me3 marks for epigenetic silencing of vial transcription (Fig 9A). The transcription activity of cccDNA, predominantly the production of pgRNA, will be significantly repressed. In case of ZNF638 deficiency in HBV infected host cells, the failure of proposed epigenetic silencing machinery on viral gene expression leads to the (re)activation of HBV life cycle, with enhanced the transcription of viral mRNAs and pgRNAs, as well as increased infectious HBV particles along with raised levels of cccDNAs (Fig 9B). ZNF638 has recently been demonstrated to interact with HUSH complex preferentially at non-coding loci of genomic DNA, and the recruitment of SETDB1 was able to confer repressive histone modification of H3K9me3^22^. Not only integrated viral DNAs, such as from retroviruse infection, be repressed through HUSH mediated gene silencing, episomal viral DNAs were also found to interact HUSH component proteins, including wild type AAVs^21, 22^. Here, we reported another case of HBV cccDNA, which existed in the form of histone decorated microchromosomes and plays a crucial role in viral persistence and antiviral therapy resistance, especially in chronic HBV infection as a high risk for terminal liver diseases.

**Fig 9.**
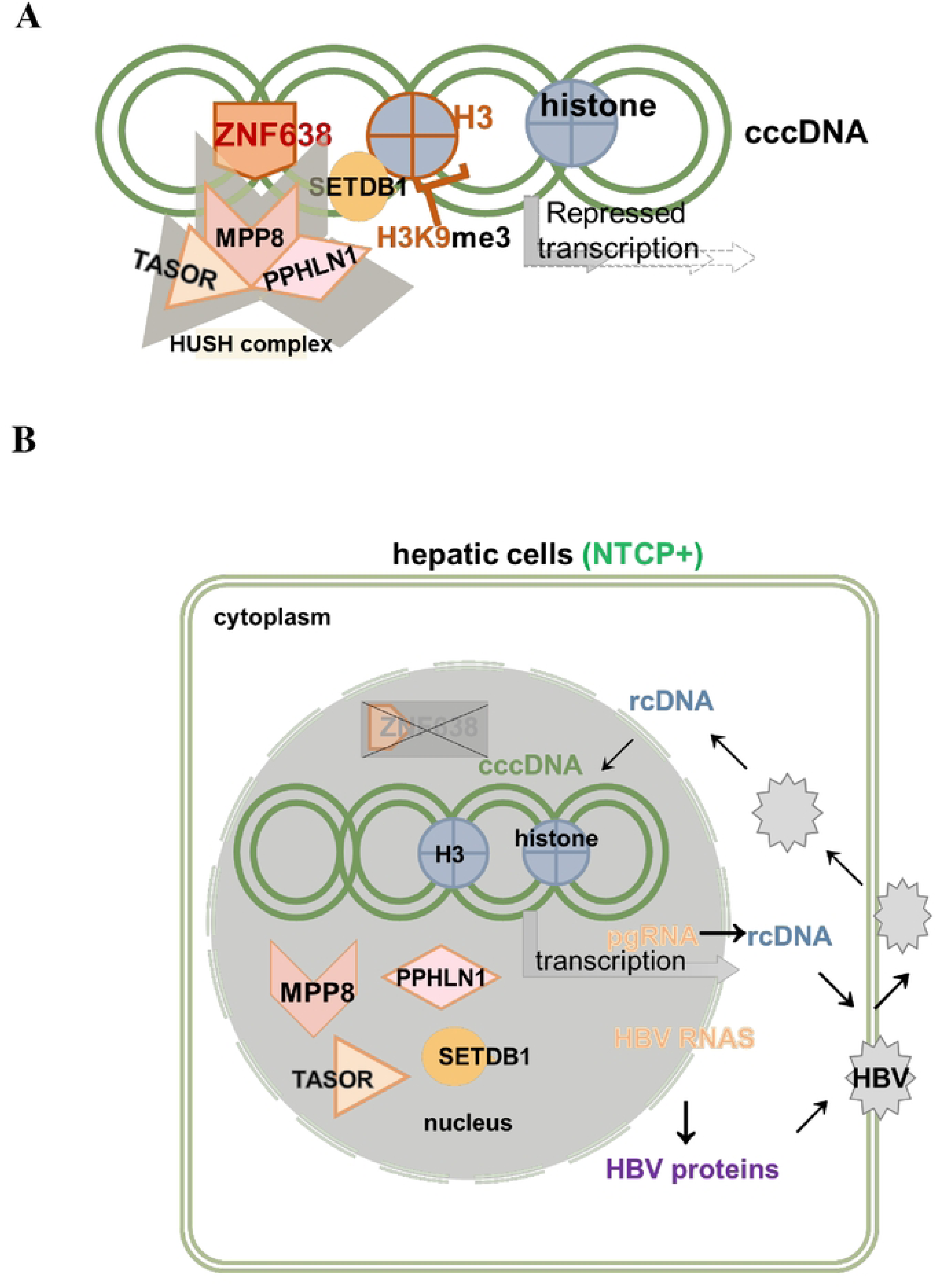
A proposed working model illustrating the role of ZNF638 to repress HBV transcription. **(A).** The direct binding of ZNF638 to cccDNA recruits the HUSH complex, enabling SETDB1 to write H3K9me3 marks onto cccDNA minichromosomes. This epigenetic silencing of viral transcription may serve as a part of the host defense mechanism in HBV infected cells. (**B**). In HBV infected ZNF638 deficient hepatocytes, without HUSH mediated SETDB1 activity and H3K9me3 dependent epigenetic silencing, the transcription of cccDNA is released at a constant high level, leading to active HBV replication, cause aggravation of pathology and compromising anti-viral therapies.

The formation and existence of cccDNA microchromosomes complicate the physiological regulation of their transcription, particularly under the impacts of various epigenetic modifications^29^, it is plausible that HUSH mediated silencing and associated machinery also apply to episomal HBV genomes, especially in their cccDNA forms. As ZNF638 was a known host factor able to repress retroviral transcription, it is worthwhile to setup host conditions using knockout or overexpressing cell models to evaluate the epigenetic regulation of HBV DNAs (Fig 1, 2). Eukaryotic cells employ epigenetic mechanisms to modulate endogenous gene expression and, in the case of invading pathogens, act as a defense mechanism by suppressing foreign genes. This status between occupancy and release at the certain targeted locus may offer adaptable regulation for controlling viral gene expression in response to various cellular and viral activities. The decoration of histones and formation into microchromosome allow HBV cccDNAs to disguise with host DNA features to avoid clearance, whereas host cells, in turn, need to develop a mechanism to repress the existing viral transcription through proper recognition. Considering the high mutation rate of HBV DNA, it is more practical to achieve silencing through histone-mediated epigenetic marking rather than by directly modifying the viral genome DNA sequence, such as through methylation of nucleotide bases.

For HUSH to exert its repressive function at the target loci, it requires certain interacting host factor to recognize the viral sequence, where ZNF638 appears to be a natural candidate. It was previously reported that ZNF638, as a member of the zinc-finger protein family, was capable of binding to the consensus sequence 5’-CCCCCG/C-3’ on DNA^30^. It was also reported that H3K9me3 modifications mediated by ZNF638 were primarily localized to the ITR regions, promoter regions, and GC-rich transgene sequences of the AAV genome^21, 22^. An important finding from the present study was that ZNF638 bound to the specific regions over *preS2* (91-318 nt) gene and *S* (445-778 nt) gene of HBV cccDNA (Fig 4), which was different from earlier reports of cytosine-rich features. Recent studies showed that integration-deficient murine leukemia virus (MLV) genomes underwent silencing by direct interaction with ZNF638, then recruited the HUSH components and the methyltransferase SETDB1 to deposit repressive H3K9me3 histone marks^22^. There is also a report suggesting that ZNF638 suppressed the expression from unintegrated genomes of HIV-1 and Mason-Pfizer monkey virus without the involvement of the HUSH complex^22^. We believe the binding of ZNF638 to HBV cccDNA is a requirement for increased histones of H3K9me3 modifications (Fig 6H, I), as opposed to other types of H3K4me3 and H3K27ac (Fig 5) for the epigenetic silencing of cccDNA transcription. Although ZNF638 possessed an inhibitory zinc finger protein, the binding of ZNF638 to HBV DNA alone appeared to be insufficient to shut down viral transcription without the participation of SETDB1 (Fig 6). This may partially explain that upregulation of ZNF638 expression as a compensatory response to HBV infection does not always lead to complete block of viral replication as observed in clinical data (Supplementary Fig 8), as well as in ZNF638oe HepAD38 or HepG2.2.15 cells. At this point, we were unable to determine whether the interaction of ZNF638 to cccDNA was solely *via* its zinc finger domains. Due to the limitation of the cell models used in the experiment, it remained not clear on whether the possible binding of ZNF638 to integrated HBV sequences was through exactly same recognition sequence patterns. In non-virus infected cells, ZNF638 was also suggested to involve DNA repair process, there was a possibility that the DNA recognition sites of ZNF638 might not have to be sequence specific short motifs as discovered in numerous ZF family transcription factors.

HUSH complex was known for its potent function to recruit and activate SETDB1 histone methylase, and we found SETDB1 played a crucial role in mediating the silence of HBV cccDNA transcription (Fig 7). We demonstrated that the function of SETDB1 to mark repressive H3K9me3 to suppress the transcription of cccDNAs with high efficiency, and the process required the participation of HUSH complex. The results showed that knockdown of each component of HUSH crippled the inhibition of on HBV transcription (Fig 7). However, the ChIP-Seq assays did not show a complete overlap between MPP8 peaks and ZNF638 peaks on HBV DNAs. We postulated that this might be because of the interaction of HUSH to cccDNAs was indirect (with required ZNF638 for DNA binding) or the local availability of the complex components varied due to their different abundance.

The silencing action of HUSH at the HBV integrated locus within the host genome also needs to be considered, given the fact that ZNF638 has been reported to interact with HUSH at sites of retrovirus integration. The binding of ZNF638 to the integrated HBV DNA in HepAD38 genome was indeed detected, however it only comprised of less than 1% of total detected ZNF638 signals in sequencing reads (Fig 4 and Supplementary Fig 6), suggesting the silencing effect of ZNF638 on the integrated HBV genome played a minor role in HBV transcription and replication, although could not be completely ignored. Recruitment of HUSH to replicated chromatin enabled inheritance of the repressed state following DNA replication, consistent with transcription being required for the maintenance of HUSH-mediated H3K9me3 at the target locus. RNA-mediated recruitment of HUSH can respond to increased transcription from its repressed locus, which may occur during cell replication. It implies that when similar mechanism functions on cccDNA microchromosomes, the repression marks could be maintained during HBV replications. However, it is noteworthy that determining whether nascent transcribed viral RNA is directly involved in the recruitment of HUSH is another important question that needs to be addressed. To this end, it is particularly significant, given that ZNF638 also contains predicted RNA-binding domains^30^. The epigenetic modifications of the HBV cccDNA microchromosomes, and their roles in regulating HBV transcription, represent a crucial area necessitating further research. In particular, factors that contribute to the deposition of repressive marks on the cccDNA, such as H3K9me3, are of significant interest. Complexes like ZNF638 and HUSH, which are involved in the ‘writing’ of these marks, emerge as key players meriting focused investigation.

In search to expand the understanding of ZNF638 with its function in clinical context of HBV infection, we explored the current available data in public domains, Analysis of a RNA-Seq dataset derived from fine needle aspirates (FNAs) of 74 CHB patients and 9 healthy donors (GSE230397) allowed us to identify the connection between the expression levels of ZNF638 and the HBV associated phenotypes. The ZNF638 expression in immunotolerant CHB group significantly increased than that of healthy control group, as well as those of immune active and HBeAg negative patient groups (Supplementary Fig 8A). This indicated that ZNF638 was associated with HBV replication, since highest levels of HBV titer was detected at immunotolerant phase (>10^7^ IU/ml, EASL 2017 guidelines), whereas low levels were maintained in immune active and HBeAg negative phase. We also presented the ZNF638 expression data of HCC tissues with HBV infection and compared to tissues from HBV negative patients in TCGA database and GSE135631 dataset. The levels of ZNF638 predicted completely different on the survival outcome, despite higher levels were general detected in HBV-related HCC individuals (Supplementary Fig 8B, C and D). Given the fact that CHB infection of its progression prone to early onset of HBV-related HCC, it once again suggested a possible role of ZNF638 in HBV infection and related host cell response. The discovered role of ZNF638 on HBV suppression was also supported by clinical statistics in chronic HBV infection and the progression of HBV-related HCC. However, the effect of ZNF638- and HUSH complex-mediated silencing in vivo remains to be further determined, especially the long-term biological effects. And it suggested that its expression could affect the effectiveness of HBV-targeted siRNA therapy (Fig 8). Future studies evaluating host-silencing mechanisms in HBV other animal models could offer additional valuable insight into HBV biology as well as provide a new direction for devising strategies for HBV therapy.

In summary, this study presents an initial discovery of ZNF638 interacting with both cccDNA and HUSH complex to suppress HBV transcription *via* a mechanism of H3K9me3 dependent epigenetic silencing. ZNF638 was identified to bind the *preS* and *S* gene regions of the HBV genome, thereby recruit the HUSH complex to facilitate SETDB1 inscribing H3K9me3 marks to silence cccDNA transcription. Our finding provides new insights into the mechanism by which HUSH silencing system modulates DNA viruses, emphasizing the importance of epigenetic silencing of cccDNA as in the histone decorated episomal forms. The results also strongly suggested that the host levels of ZNF638, as a potent host limiting factors of viral genome epigenetic modulation activities, needs to be concerned in future anti-HBV therapy.

## Materials and Methods

### Plasmids, siRNAs and reagents

ZNF638 expressing plasmids were constructed by amplification of the complete CDS (coding sequence) of ZNF638 from human liver cancer cell line HepG2, using high fidelity polymerase (Takara, Beijing, China) and cloned into piggyBac Dual promoter vector. The prcccDNA/pCMV-Cre recombinant plasmid system was a kind gift from Professor Qiang Deng of Fudan University^24^. The designed siRNAs targeting to ZNF638, MPP8, PPHLN1, TASOR and SETDB1 (sequences listed in Supplementary Table S1) were synthesized from General Biol (Anhui, China). Rabbit IgG and Mouse IgG were products from Proteintech (Rosemont, IL, USA). Tetracycline was purchased from Sigma-Aldrich (Saint Louis, MO, USA).

### Preparation of HBV infectant from patient serum

Four serum samples from HBV-infected HCC patients from Beijing You-An Hospital, Capital Medical University (Beijing, China) were used in this study. The experiment was approved by the Ethics Committee of Beijing You-An Hospital (7222096). Clinicopathological information from medical records of involved patients were summarized in Table S2. A discontinuous sucrose density gradients were prepared using 1ⅹPBS, pH 7.2. Adding carefully the serum to centrifuge tubes containing prepared sucrose gradient, centrifugation was performed at 4°C 140,000 g for 8 h using L-80XP high-speed centrifuge. Remove the supernatant, and then add in 10% FBS (Biological Industries, Israel) containing PBS to resuspend the pellet.

### HBV infection

As described^31^, the media from HepAD38 cells were collected in every 3 days during a 4-10 day induced culture in tetracycline free conditions. The obtained media were cleared through a 0.45 μm filter and precipitated with 40% PEG 8000. The precipitates were washed and resuspended with FBS. The amount of HBV DNA was quantified by real-time PCR and the primer sequences are listed in Table S3. HepG2-NTCP cells were constructed as previously described^32^, which were cultured in DMEM supplied with 10% FBS. Native HepG2-NTCP cells, or transfected HepG2-NTCP cells with Lipofectamine 3000 Transfection Reagent (Invitrogen, Carlsbad, CA, USA) were exposed to prepared HBVs in the presence of 4% PEG 8000 and 1.5% DMSO for 20 h. The infected cells were then washed with PBS and maintained in DMEM with medium change in every 2-3 days.

### Knockout for ZNF638 deficient cells

As previously described^33^, a circular RNA encoding Cas9 protein was designed and constructed. Briefly, Cas9 mRNA, group I self-splicing intron, and IRES sequences were chemically synthesized and cloned into a PCR-linearized plasmid vector containing a T7 RNA polymerase promoter by General Biol. *In vitro* transcription was carried out with TranscriptAid T7 High Yield Transcription Kit (Thermo Fisher Scientific, Waltham, MA, USA) as recommended by the manufacturer using a primer specific for a region internal to the Cas9 circRNA. For splice junction sequencing, splicing reactions enriched for circRNA with RNase R (Geneseed, Guangzhou, China) and then column purified were heated at 65 °C for 5 min and cooled on ice for 3 min to standardize secondary structure. 100 ng of ZNF638 sgRNA vector was reverse transfected alone or cotransfected with 150 ng of RNase R-treated Cas9 circRNA into 50,000 HepAD38, HepG2.2.15 or HepG2-NTCP (gifts from professor Dexi Chen) cells/500 uL per well of a 24-well plate using Lipofectamine 3000. After transfection 5-7 days, the sgZNF638 in cells was verified by qPCR, western blot and sequencing (Supplementary Fig 1), and select a cell pool for stable passage.

### Fluorescence in situ hybridization (FISH) and immunofluorescence (IF)

HepG2-NTCP cells were infected with 6×10^5^ copies/ml HBV and cultured on 24-well plates. In FISH assays, the cells were subjected to pretreatment with either RNase A (Vazyme) for HBV DNA detection, or RNase A and T5 exonuclease (Beyotime Biotechnology) for detecting cccDNA. The cells were then fixed with 4% paraformaldehyde (PFA) for 10 min, permeabilized with 0.5% (w/v) Triton X-100 for 15 min. Following blocking at room temperature in pre-hybridization solution (2×SSC and 10% formamide) for 30 min, the cells were hybridized with probes overnight at 37 °C. The sequence of Cy3-labeled HBV DNA probes (from General Biol) was complementary to the conserved segment of HBV DNA (NC_003977.2) from nt. 1091 to 1557. Followed washing three times with 2 × SSC, the samples were subjected to IF staining. The cells were refixed in 4% paraformaldehyde for 20 min and blocked (2% BSA). The cells were incubated subsequently with a primary antibody targeting ZNF638 (Bethyl, Texas, USA) for 2 h at room temperature, and a CoraLite488-conjugated secondary antibody (Proteintech) for 1 h in the dark. The nuclei were stained with DAPI (Solarbio, Beijing, China) prior to imaging under a confocal system (TCS SP8 STED, Leica Microsystems, Germany). Images were acquired and processed using Leica software (Leica). The background level of non-specific binding was determined from control samples prepared without the addition of the primary antibody.

### Chromatin immunoprecipitation and ChIP-seq

Chromatin immunoprecipitation (ChIP) experiments were carried out as described with minor modifications^34^. Briefly, the cells were fixed with 1% formaldehyde at 37 °C, then washed with pre-cooled PBS for 3 times. The collected cells were lysed in nuclear lysis buffer (1% SDS, 50 mM Tris-HCl, 5 mM EDTA) and sheared by sonication to generate DNA fragments. After centrifugation, the diluted supernatant was precleared with protein A/G magnetic beads (Gene-Protein Link, Beijing, China) in dilution buffer (1% TritonX-100, 2 mM EDTA, 150 mM NaCl, 20 mM Tris-HCl). Immunoprecipitation was performed at 4°C overnight by incubating with desire antibody, followed by incubation with protein A/G magnetic beads for additional 2 h. Samples were washed and digested with proteinase K at 65 °C for 10 hours. The immunoprecipitated proteins were subjected to western blot and quantified by densitometry for normalization. The retrieved DNA was assayed by both conventional PCR and qPCR and the results were normalized to levels of input DNAs and presented in relative fold enrichment compared over the control, normalized to input DNA using the ΔCt method, the calculation was ΔCt = Ct(input) - Ct(immunoprecipitation), and the result was expressed as a percentage of the input.

The ChIP-seq library was prepared using VAHTS Universal Pro DNA Library Prep Kit for Illumina (Vazyme, Nanjing, China). Libraries were purified, quantified, multiplexed and sequenced with 2× 50-bp pair-end reads on Illumina Novaseq platform (Illumina, NovaSeq X Series specifications). A serie of software packages was deployed for bioinformatic analyses, including FastQC (Babraham Bioinformatics) (v0.11.7) cutadapt37 (v1.16), HISAT2 (v2.1.0), SAMtools (v1.9), and ChIPseeker (v1.26.2).

### HBV cccDNA-ChIP

HBV-producing HepAD38 cells cultured in Tet-free medium and in HepG2-NTCP cells infected for 6 days, HBV cccDNA-ChIP experiments were conducted following a modified ChIP protocol (34). In brief, cells were fixed with formaldehyde and suspended in nuclear lysis buffer. The samples were subjected to ultrasonication at a selected condition for preserving the integrity of cccDNAs. Subsequently, the fragmented DNAs were digested with T5 exonuclease (Beyotime Biotechnology) at 37°C for 1 hour to removed other non-cccDNA forms of HBV DNAs. Specific antibodies of xx,xx or xx were used for chromatin immunoprecipitated along with protein A/G agarose beads. The extracted DNAs were quantified by qPCR using primers specific to cccDNAs.

### Assay for HBV replication

Secreted HBeAg in infected cell cultures were assayed using a Human Hepatitis B E Antigen (HBeAg) ELISA kit (MSK, China) with the recommended protocol following the manufacturer’s instructions. The HBV DNA was extracted using the Alkaline lysis method^35^ from the medium or animal serum. Following incubation at 37 °C for 30 minutes with 0.2 M NaOH solution, 0.2 M HCl was added to neutralize the reaction. Centrifugation at 12,000 rpm was performed to remove excess NaCl. The obtained HBV-DNA was quantified by qPCR using a Hepatitis B Viral DNA quantitative fluorescence diagnostic kit (Sansure Biotech, Changsha, China). Total RNA was isolated from infected cells or liver tissues of mice and used for the preparation of cDNAs. The HBV-RNA was quantified by RT-qPCR. The primer sequences were listed in Table S3.

### HBV cccDNA isolation

DNA extraction was performed following a modified Hirt extraction procedure^36^. Briefly, the cells were lysed on ice for 30 min in lysis buffer A (50 mM Tris-HCl pH 7.4, 1 mM EDTA, 1% NP-40) containing a complete protease inhibitor cocktail. The pelleted nuclei from centrifugation were resuspended in lysis buffer B (10 mM Tris-HCl, 10 mM EDTA, 150 mM NaCl, 0.5% SDS, Proteinase K 0.5 mg/ml) and incubated overnight at 37 °C. Phenol-chloroform (1:1) was used for DNA extraction, and the purified products were precipitated in ethanol. The obtained protein-free DNA was treated with DNase-free RNase and digested with a restriction enzyme (*Eco*R I) before quantitation. The extracted cccDNAs were treated at 37°C with T5 exonuclease for 1 h for the removal of other contaminant forms (including relaxed circular DNA (rcDNA), single-stranded DNA (ssDNA) or linear double-strand DNA (dsDNA)). The final isolated cccDNAs were quantified by qPCR in 20 μl reaction volume, using 2 μl digested cccDNA by specific primers for cccDNA (targeted across the single-stranded (SS) gap region of rcDNA), as listed in Table S3.

### Nuclear cccDNA pulldown

A biotin tagged 49-mer oligonucleotide complementary to the conserved sequence of HBV cccDNA (nt: 1865 to 1913; NCBI: NC_003977.2) was synthesized by General Biol, giving rise to the cccDNA oligonucleotide probes. HepAD38 cells were cultured in Tet-free medium to induce HBV replication for 6 days. HepG2-NTCP cells were infected with 6×10^5^ copies/ml HBV for 6 days to accumulate cccDNA. The cells were collected, and the nuclear fractions were isolated using a nuclear extraction kit (Cayman Chemical, Michigan, USA). The nuclear fractions were treated with T5 exonuclease at 37 °C for 1 h to remove the HBV DNA in rcDNA, ssDNA and dsDNA forms. The obtained cccDNAs were probed with 1.25 nmol of biotinylated cccDNA oligonucleotide bait at 4°C for 16 h using a gentle rotator. Next, 50 ml of streptavidin coated beads (Beyotime Biotechnology, China) was added to the reaction and incubated for 4 h at 4°C under constant slow rotation. The beads were precipitated and washed in wash buffer containing 5 mM Tris-HCl (pH 7.5), 0.5 mM EDTA, 1 M NaCl and 0.05% Tween-20 3 times. The cccDNA-bound proteins were eluted from the beads with sodium dodecyl sulfate (SDS) sample loading buffer, resolved by SDS-PAGE and subjected to Western blot analysis with antibodies.

### Southern blotting

Southern blotting was performed as described^36^. Briefly, the extracted HBV cccDNA sample was subjected to 1.2% agarose gel electrophoresis and transferred onto an Hybond-N+ membrane (Beyotime Biotechnology). The Hybond-N+ membrane was cross-linked in a UV cross-linker chamber with a UV energy dosage of 25 J, followed by probing with DIG-labeled probes (Absin, China) for 24 hours at 42°C. The membrane was then blocked and incubated with Anti-Digoxigenin-AP (Absin) for 1 hour at 37°C. After washing for 15 minutes, add 10 ml of freshly prepared NBT/BCIP staining solution (Absin), place it in a dark room or dark box, and let it develop color for 0.5 to 16 hours.

### *In vivo* experiments using HBV transgenetic mice

The C57B/6N-Tg(1.28HBV)/Vst mice were purchased from Beijing Vitalstar Biotechnology Co.,Ltd (China). and maintained under pathogen free condition. GalNac-siHBV (3 μg/g BW) and GalNac-siZNF638 (3 μg/g BW) were administrated through subcutaneous injection at day 0. Blood samples were collected from the eye socket at day 8, 15, 22 and 29. After 29 days of maintaining, mice were sacrificed and liver tissues were collected. For histological analysis, mice liver tissues from autopsy were fixed 10% formalin for 48 h before delivered to Beijing Tongnong Testing Technology Co., Ltd for further processing, sections and staining for HbeAg and ZNF638, besides H&E.

### RNA extraction and RT-qPCR

Total RNA was isolated using RNA-Quick Purification kit (Yishan biotechnology Co. LTD, China). HiScript II Q RT Kit (Vazyme, China) was used for reverse transcription. AceQ qPCR SYBR Green Master Mix (Vazyme) was used to quantify gene expression level from the obtained cDNA. The primers for detecting are listed in Table S3. GAPDH was used as the loading reference. The quality and concentration of cDNAs were evaluated using a Quantitative Real-time PCR system (Archimed X6, Rocgene, China).

### Western blotting

Western blotting was performed as described^36^. Briefly, cell lysates or isolated HBV cccDNA samples were subjected to 10% SDS-polyarylamide gel electrophoresis (SDS-PAGE), then transferred onto polyvinylidene fluoride (PVDF) filters. The probing antibodies against the following antigens: ZNF638 (Bethyl, USA), MPP8 (Proteintech), H3K9me3 (Abcam, UK), H3K27ac (Gene Tex, San Antonio, USA), H3K4me3 (Active motif, Carlsbad, CA, USA), H3 (Cell Signaling Technology, Massachusetts USA), and GAPDH (Proteintech) were used for overnight incubation at 4 °C following the blocking procedure. Goat anti-Rabbit secondary antibody, Goat anti-Mouse secondary antibody with conjugated HRP were purchased from Proteintech and used for developing chemluminescent signals by Minichemi chemiluminescence imager (Beijing sage creation).

### Statistical Analysis

All statistical analyses were performed using GraphPad Prism V8. Unpaired Student’s t-test and one-way ANOVA were applied to determine the statistical significance of differences of different data groups. Data were presented as mean±standard deviation (SD). (* *p* < 0.05, ** *p* < 0.01, *** *p* < 0.001, NS: no significance).

## Data availability

Raw data from the ChIP-seq experiments have been deposited to the GEO database (https://www.ncbi.nlm.nih.gov/geo/) under the accession number GSE262325.

## Authors’ contributions

S.C. and W.D. designed the research. S.F.M., X.W.S, G.Y.Z, J.W., S.Y. and F.L performed the experiments. S.F.M., X.W.S, G.Y.Z, B.W.D., J.W., S.Y., F.L, Q.W., S.C. and W.D. analyzed the data. S.F.M. and S.C. wrote the paper.

## Ethics approval and consent to participate

Serum samples utilized in the study were collected within protocols approved by the Ethics Committee of Beijing You’an Hospital, with the approval number 7222096.

## Acknowledgements

This work was supported by the National Natural Science Foundation of China (No. 82073215 to S.C.), the Beijing Natural Science Foundation Program (No.7222001 to W.D.). We would like to thank Prof. Dexi Chen for providing human hepatocellular cell lines, Prof. Binwei Duan for serum samples, Prof. Qiang Deng for prcccDNA/pCMV-Cre recombinant plasmid system, Prof. Fengming Lu for helpful discussions.

## Conflict of Interest

The authors declare no conflict of interest.

## Notes

### Competing Interest Statement

The authors have declared no competing interest.

